# Repression of the Wnt pathway effector TCF7L2 reverses lethal cachexia in mice with intestinal cancers

**DOI:** 10.1101/2024.11.22.624913

**Authors:** Mei Ling Leong, Christiane Ruedl, Klaus Karjalainen

**Affiliations:** School of Biological Sciences, Nanyang Technological University

**Keywords:** TCF7L2, Wnt signaling, colon cancer, small intestine cancer, cachexia, muscle atrophy

## Abstract

Hyper-activation of the canonical Wnt signaling pathway drives small intestine and colon tumors. As the major Wnt pathway effector in healthy intestines, TCF7L2 is a suspected oncogene in both cancer types. However, this has been challenging to verify because *Tcf7l2* knockout is lethal. To circumvent lethality, we generated a novel transgenic mouse that allows dose-dependent, systemic, inducible and reversible repression of endogenous *Tcf7l2* expression. Using this mouse, we demonstrate that TCF7L2 is essential for early adenoma development in the small intestine (*Apc*^Min/+^ mouse model) but not colon (DSS-treated *Apc*^Min/+^ mouse model). Once established however, neither small intestine nor colon adenomas require TCF7L2 for maintenance. Despite this, *Tcf7l2* repression rescues both types of cancer mice from lethal cachexia—a prevalent cancer comorbidity characterized by debilitating weight loss and skeletal muscle atrophy. In colon cancer cachexia, elevated TCF7L2 in the gastrocnemius muscle induces atrophy by activating the transcription of multiple atrophy genes within the ubiquitin-proteasome and autophagy-lysosome systems. Hence, repressing *Tcf7l2* normalizes atrophy gene expression back to non-cachectic expression levels and restores muscle mass. The cachexia recovery mechanism in small intestine cancer remains undefined but is independent of the gastrocnemius. This study shows that systemic and partial *Tcf7l2* repression is both well-tolerated and effective in rescuing moribund cancer mice from cachexia-induced death. Hence this is a promising treatment strategy for cancer patients suffering from cachexia. Additionally, our transgenic mouse is a valuable tool to study muscle atrophy across other conditions including aging, diabetes and neuromuscular disease.

## Introduction

The canonical Wnt signaling pathway is involved in diverse biological processes including cell proliferation, stem cell maintenance and tumor pathology, among others (1, 2). When the pathway is activated, β-catenin is stabilized, and it interacts with one of the four TCF/LEF transcription factors to transcribe Wnt target genes. As the most abundant TCF/LEF protein in healthy colon and small intestine, TCF7L2 is the major transducer of Wnt signal in both organs. It is essential for maintaining *Lgr5*^+^ intestinal epithelial stem cells (3) and ensuring homeostatic turnover of the intestinal epithelium (3, 4). Since *Lgr5*^+^ epithelial stem cells are the origin of both small intestine and colon tumors (5), the ability to maintain these cells implies that TCF7L2 is an oncogene.

However, contradictory roles for TCF7L2 in colon tumors have arisen from several studies using different strategies to target the gene (6–10). TCF7L2’s role in small intestine tumors is poorly understood because this cancer type is less well-studied.

TCF7L2 may also be involved in cachexia, which is a hypermetabolic state that affects 40-80% of cancer patients (11) and is more prevalent in gastrointestinal cancers (12, 13). This is because TCF7L2 maintains metabolic homeostasis in healthy individuals by regulating various functions in metabolic organs such as white adipose tissue, pancreas, and liver (14–16). Cachexia is induced by tumor-derived cachectic factors, including growth factors, cytokines and metabolites (17, 18), which act systemically on multiple distant organs to promote decreased appetite, skeletal muscle atrophy and adipolysis(19, 20). The loss of muscle mass during early and mild cachexia is modest (12). However, as cachexia progresses into the late and severe refractory stage, muscle atrophy becomes so drastic (12) that it may cause death by cardiac or respiratory arrest (19). A recent phase 1b clinical trial has shown promise in restoring appetite and reversing cachexia in cancer patients (21) by blocking the interaction of cachectic factor GDF-15 with its cognate receptor GFRAL on brain stem neurons (22–25). Apart from this, other proposed treatments for cachexia such as supplementing nutritional requirements, inhibiting muscle atrophy and suppressing pro-inflammatory cytokines, are minimally effective (26). Therefore, better treatments for cachexia are required to improve the quality of life and extend the survival of cancer patients. With this in mind, we aimed to investigate the potential roles of TCF7L2 not only in the local context of colon and small intestine tumors, but also in the underlying systemic cachexia.

## Results

### Robust regulation of endogenous *Tcf7l2* expression in transgenic mouse

As the canonical Wnt signaling pathway is crucial for many homeostatic processes across different organs and TCF7L2 is ubiquitously expressed in many of these organs, it is important to ensure that perturbing this gene would not be detrimental to the mouse host. Hence, we leveraged the Tet-inducible system to achieve an effect similar to administering a therapeutic drug—partial yet ubiquitous *Tcf7l2* repression. This newly generated double transgenic mouse strain (Fig. 1a) had the tetracycline repressor (rTetR-KRAB) and tetracycline response element (TRE) inserted into the *Rosa26* and *Tcf7l2* loci, respectively (Supplementary Fig. S1). Feeding mice with doxycycline-containing food pellets rapidly and completely repressed endogenous *Tcf7l2* mRNA and protein expression in the small intestine within three days (Fig. 1b-d). Repression was reversible since both mRNA and protein expression returned to their original levels upon withdrawal of doxycycline treatment (Fig. 1b-d). When *Tcf7l2* was continuously repressed, mice became moribund within 10 days due to the loss of the small intestine epithelium (Fig. 1e-f). This is identical to the lethal phenotype of other germline (3, 4) and adult (6) *Tcf7l2* knockout mice, thus validating our transgenic mouse.

**Figure 1.**
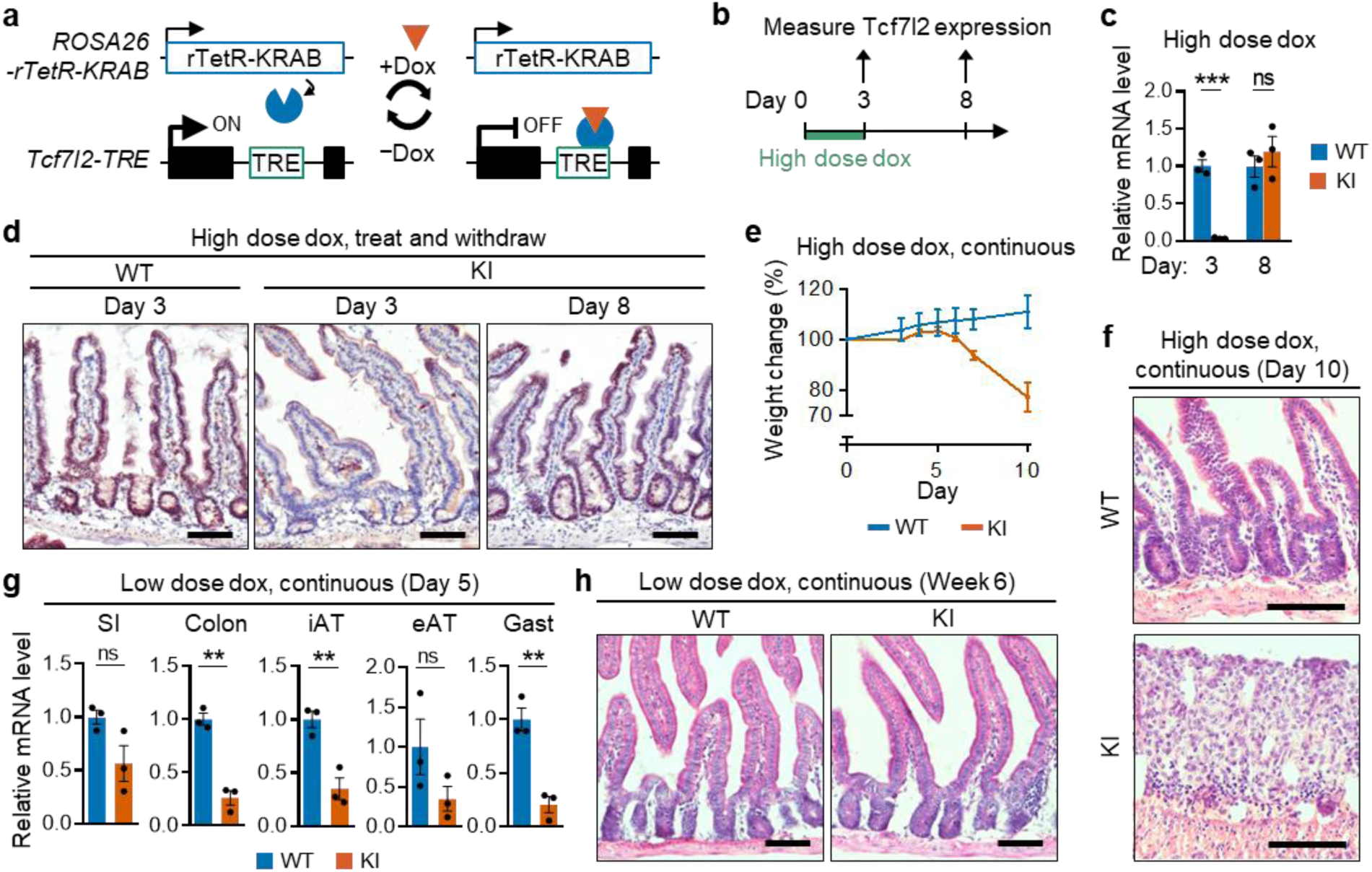
Robust regulation of endogenous TCF7L2 expression in transgenic mice. **(a)** Schematic representation of the Tet-inducible system to regulate endogenous *Tcf7l2* expression in double transgenic mouse. The reverse tetracycline trans-silencer (rTetR-KRAB) is constitutively expressed from the ROSA26 genomic region. Treatment with doxycycline (Dox) causes rTetR-KRAB to bind tetracycline response element (TRE) and transcriptionally repress *Tcf7l2*. *Tcf7l2* expression is restored upon cessation of doxycycline treatment. **(b-h)** WT (*Rosa26*^rTetR-KRAB/WT^*Tcf7l2*^WT/WT^) and KI (*Rosa26*^rTetR-KRAB/WT^*Tcf7l2*^TRE/TRE^) mice were treated with either high or low dose doxycycline (dox) for various time durations. **(b)** Experimental timeline for high dose doxycycline treatment and withdrawal. *Tcf7l2* **(c)** mRNA and **(d)** protein expression in the small intestine following doxycycline treatment and withdrawal as outlined in (b). Data are mean ± SEM. Statistical significance was assessed by Student’s t-test. All experimental groups *n* = 3. ***P < 0.001 and ns, not significant. Scale bar 50μm. **(e-f)** Mice were continuously treated with high dose doxycycline for ten days. **(e)** Body weight with respect to day 0 and **(f)** H&E staining of small intestine on day 10. Data are mean ± SEM. WT *n* = 3 and KI *n* = 6. Scale bar 50μm. **(g-h)** To partially repress *Tcf7l2*, mice were continuously treated with low dose. **(g)** Relative *Tcf7l2* mRNA expression in the indicted tissues on day 5 and **(h)** H&E staining of small intestine on week 6. Data are mean ± SEM. Statistical significance was assessed by Student’s t-test. All experimental groups *n* = 3. **P < 0.01 and ns, not significant. Scale bar 50μm. eAT, epididymal adipose tissue. iAT, inguinal adipose tissue. SI, small intestine.

Lethality can be avoided by down-titrating doxycycline dose, to achieve a partial 50% repression in the small intestine (Fig. 1g). At this level of repression, the small intestine epithelium remained intact (Fig. 1h) and mice were healthy even after one year (data not shown). Unexpectedly, low-dose doxycycline led to varying levels of *Tcf7l2* repression across different tissues. Repression reached 90% in the colon, 80% in the gastrocnemius, 95-98% in colon tumors and 60-80% in small intestine tumors (specific details on the repression levels are provided in the subsequent sections). Therefore, partial *Tcf7l2* repression is well-tolerated, yet high levels of repression can be achieved in tumors and muscles, providing us with an ideal strain to study the long-term effects of *Tcf7l2* repression on tumors and the associated cancer cachexia.

### *Tcf7l2* repression rescues moribund colon cancer mice

To model colon tumors with hyperactivated canonical Wnt signaling pathway, we mated our transgenic mouse with the *Apc*^Min/+^ strain and treated progeny mice with dextran sodium sulphate (DSS) (27). Progeny with various combinations of wildtype or transgenic alleles at the *Rosa26*, *Tcf7l2* and *Apc* loci were used either as experimental or control mice (Fig. 2a). Following DSS treatment, Col-WT and Col-KI mice grew more slowly than noCol-WT and noCol-KI mice (Fig. 2b-c), due to developing colon tumors. A sudden drop in body weight occurred 25-40 days after DSS treatment (Fig. 2c) and was associated with greatly reduced gastrocnemius muscle, inguinal adipose and epididymal adipose masses (Fig. 2d), which suggests the onset of severe refractory cachexia. Beyond this point, all Col-WT mice quickly became moribund within 2 days and were euthanized because of a 20% body weight drop (Fig. 2c). In stark contrast, Col-KI mice were rescued by initiating doxycycline treatment at the first sign of weight drop. There was a rapid rebound in body weight, muscle mass and adipose mass within 5 days (Fig. 2c-d), and survival was extended for another four to ten weeks (Fig. 2e). Continuous *Tcf7l2* repression was necessary for survival, since cessation of doxycycline treatment on day 14 led to deterioration and death (Fig. 2e). This deterioration was more gradual than in Col-WT mice due to doxycycline leaching from accumulated depots in bones (28) and adipose (29). Taken together, these findings suggest that continuous *Tcf7l2* repression rescues colon cancer mice, likely from cachexia-induced death, and significantly prolongs their lifespan. However, mice eventually die due to other complications that arise from cancer progression.

**Figure 2.**
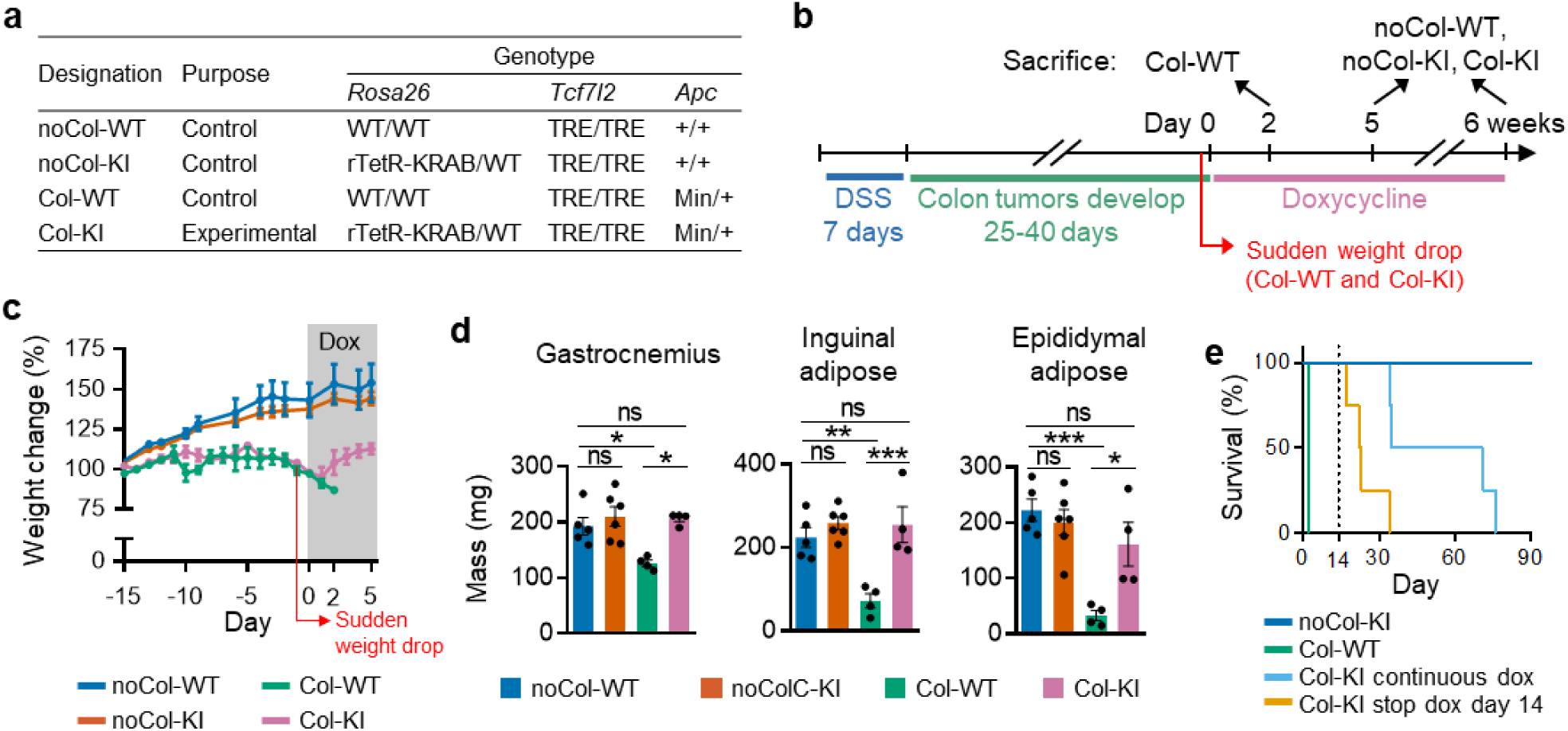
*Tcf7l2* repression rescues colon cancer mice from fatal cachexia. **(a)** Genotypes of control and experimental mice at the three loci indicated. **(b)** Experimental timeline for colon tumor development and doxycycline treatment. **(c)** Body weight with respect to day -15. **(d)** Mass of the indicated tissues from mice undergoing two days (Col-WT) or five days (noCol-WT, noCol-KI and Col-KI) of doxycycline treatment. **(e)** Survival analysis of mice given continuous doxycycline treatment (continuous dox) or when doxycycline treatment ceased on day 14 (stop dox). Data are mean ± SEM. Statistical significance was assessed one-way ANOVA with Bonferroni test. noCol-WT *n* = 5, noCol-KI *n* = 6, Col-WT *n* = 4, Col-KI *n* = 4. All experimental groups n = 4. *P < 0.05; **P < 0.01; ***P < 0.001 and ns, not significant. DSS, dextran sodium sulphate.

### TCF7L2 is dispensable for the maintenance and progression of colon tumors

Another study has shown that mouse colon tumors can be eradicated by suppressing hyperactivated canonical Wnt signaling pathway for four to eight weeks (30). The authors achieved this by restoring the expression of APC, which acts upstream of TCF/LEF transcription factors to negatively regulate the Wnt signaling pathway. Hence, we hypothesized that repressing *Tcf7l2* in Col-KI tumors would cause tumor regression because it attenuates the same signaling pathway.

Doxycycline treatment repressed *Tcf7l2* expression in Col-KI tumors by 95-98% and diminished Wnt signaling activity, as measured by the decreased expression of Wnt target genes *Lgr5* and *Axin2* (Fig. 3a). However, both genes were still slightly more elevated in Col-KI tumors compared to their adjacent normal tissues, which suggests that the Wnt signaling pathway remained slightly hyperactivated. Nonetheless, we maintained *Tcf7l2* repression for six weeks and found that colon tumors persisted (Fig. 3b-c). In fact, the proportion of large tumors in APC-KI mice increased (Fig. 3c-d), likely because small tumors continued to grow and coalesced into larger units. Albeit growth might have been stunted since tumors were smaller than expected for six weeks of continued growth. Histological analysis revealed that although most of the colonic lesions were adenomas, some had progressed to adenocarcinomas with local invasion into the submucosa (Fig. 3e).

**Figure 3.**
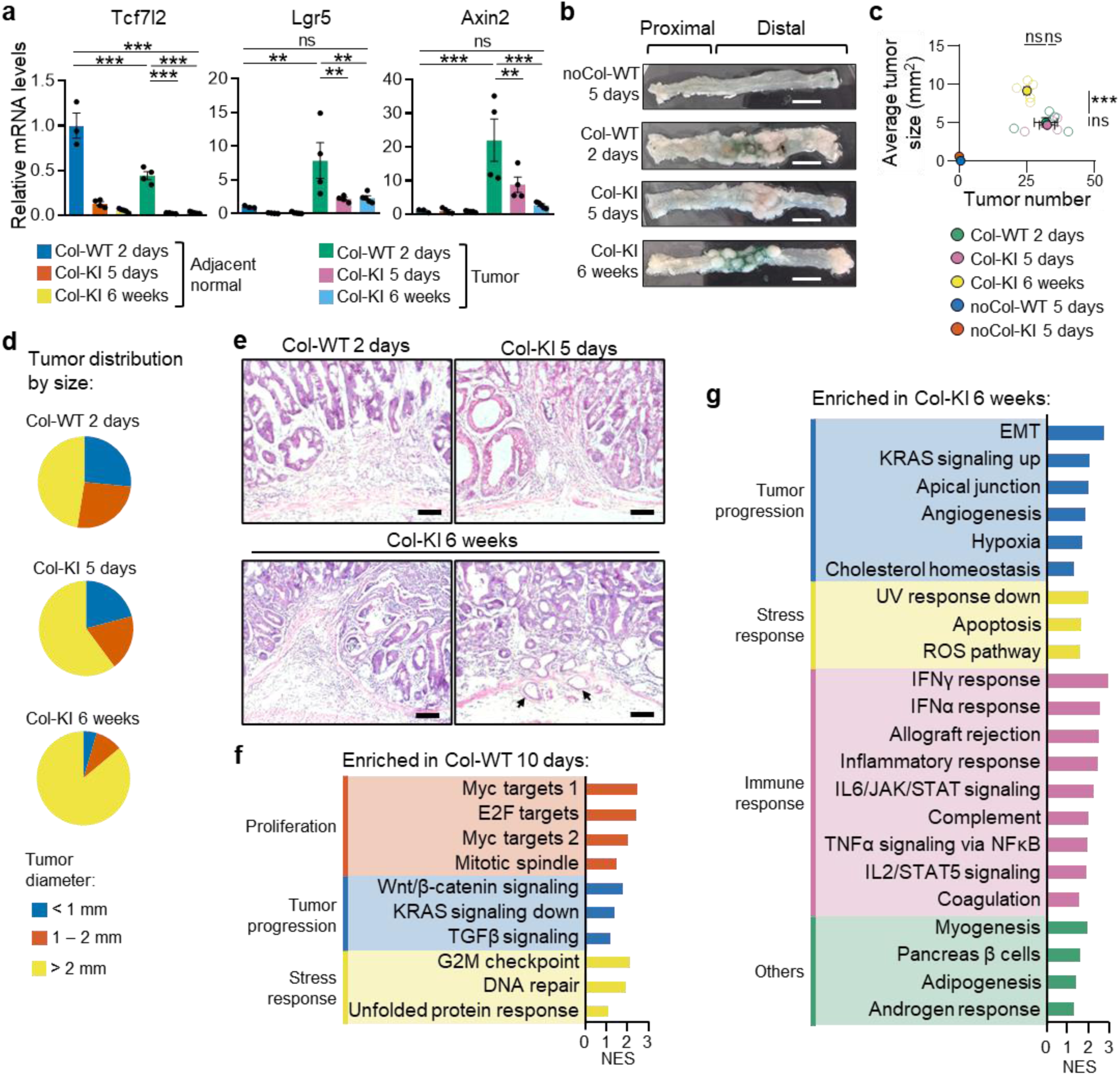
Colon tumors continue to progress despite recovery in mice. Mice were treated with doxycycline for various lengths of time, as indicated. **(a)** Relative mRNA expression of Tcf7l2 and Wnt target genes in tumors and the adjacent normal tissue. **(b)** Representative images of longitudinally opened colon. Scale bar 1cm. **(c)** Quantification of tumor burden and **(d)** distribution of tumors by size. **(e)** H&E staining of tumors. Arrows indicate adenocarcinoma growth that has invaded the submucosa. Scale bar 100μm. **(f-g)** Gene Set Enrichment Analysis (GSEA) based on bulk RNA-Seq of tumors. Doxycycline treatment was initiated several days early so that mice were not yet moribund at the point of euthanasia. Shown are Hallmark gene sets with false discovery rate < 0.25. Bars are colored according to biological function of the gene set, as indicated. NES, normalized enrichment score. Data are mean ± SEM. Statistical significance was assessed by one-way ANOVA with Bonferroni test. APC-WT 2 days (a) *n* = 3 and (c) *n* = 4, APC-KI 5 days *n* = 4, APC-KI 6 weeks *n* = 5. **P < 0.01; ***P < 0.001 and ns, not significant.

To understand how Col-WT and Col-KI tumors differ at the molecular level, we performed bulk RNA-sequencing (RNA-Seq). Col-WT and Col-KI mice were treated with doxycycline for ten days and six weeks, respectively. Here, doxycycline treatment was initiated several days before the onset of severe cachexia, so that Col-WT mice were not yet moribund at the time of euthanasia.

Differentially enriched gene sets between the two groups were identified via Gene Set Enrichment Analysis (GSEA) and categorized according to their biological process. Col-WT tumors were enriched for Myc targets, E2F targets, G2/M checkpoint, DNA repair and mitotic spindle gene sets (Fig. 3f), which are associated with proliferation and cell cycle progression (31). This indicates that emerging Col-WT tumors (day 10) were more proliferative than established Col-KI tumors (week 6). In comparison, Col-KI tumors were enriched for gene sets associated with tumor invasion and aggressiveness (32, 33), including EMT (epithelial mesenchymal transition), apical junction, angiogenesis, hypoxia and KRAS signaling (Fig. 3g). These findings might suggest that TCF7L2 is involved in promoting tumor proliferation at the expense of invasion. However, they may simply reflect differences in tumor stage since it was impossible to obtain age-matched tumors. TCF7L2 is obviously dispensable for maintenance and progression of colon tumors, because Col-KI adenomas progressed into more advanced adenocarcinomas despite prolonged *Tcf7l2* repression. Thus, the recovery of Col-KI mice was not due to changes in tumor burden.

Intriguingly, *Tcf7l2* expression was halved, yet Wnt target genes *Lgr5* and *Axin2* were highly elevated in Col-WT tumors compared with their adjacent normal tissues (Fig. 3a). This implies that Wnt signal was more highly transduced despite reduced TCF7L2 expression. It has been shown that expression of TCF7L2 is reduced while TCF7 and LEF1 are elevated in colon tumors compared to the adjacent normal tissues of colorectal cancer patients (34). Similarly, we found that *Tcf7* and *Lef1* expression were substantially elevated in tumors (Supplementary Fig. S2). This demonstrates that, similar to colorectal cancer patients, colon tumors in DSS-treated *Apc*^min/+^ mice also have a switch in TCF/LEF protein usage, with TCF7 and LEF1 becoming the predominant paralogs.

To determine whether TCF7L2 might be important in early tumourigenesis, we initiated doxycycline treatment 10 days after DSS treatment (Supplementary Fig. S3a-b), but before adenomas develop (27). This had no significant impact on tumor burden in Col-KI mice (Supplementary Fig. S3c-f), which suggests that TCF7L2 is not essential for early colon tumors. However, we cannot exclude the possibility that *Tcf7l2* repression would prevent tumor initiation because the first mutagenic events occur soon after DSS treatment (27). As mice were recovering from DSS-induced inflammation, they could not tolerate simultaneous *Tcf7l2* repression (data not shown).

### Muscle atrophy in cachectic muscles is reversed by *Tcf7l2* repression

The swiftness in the recovery of gastrocnemius muscle, inguinal adipose and epididymal adipose masses within five days of *Tcf7l2* repression in Col-KI mice (Fig. 2d), suggests that TCF7L2 plays a direct role in these tissues. Indeed, *Tcf7l2* expression in these tissues correlated with cachectic status—*Tcf7l2* was up-regulated in all three tissues of Col-WT mice during severe cachexia, as opposed to non-cachectic noCol-WT and noCol-KI, and recovered Col-KI mice (Fig. 4a and Supplementary Fig. S4). Since cachexia is strictly defined by skeletal muscle atrophy, but not necessarily adipolysis (12), we focused on understanding the link between TCF7L2 and skeletal muscle atrophy in the gastrocnemius.

**Figure 4.**
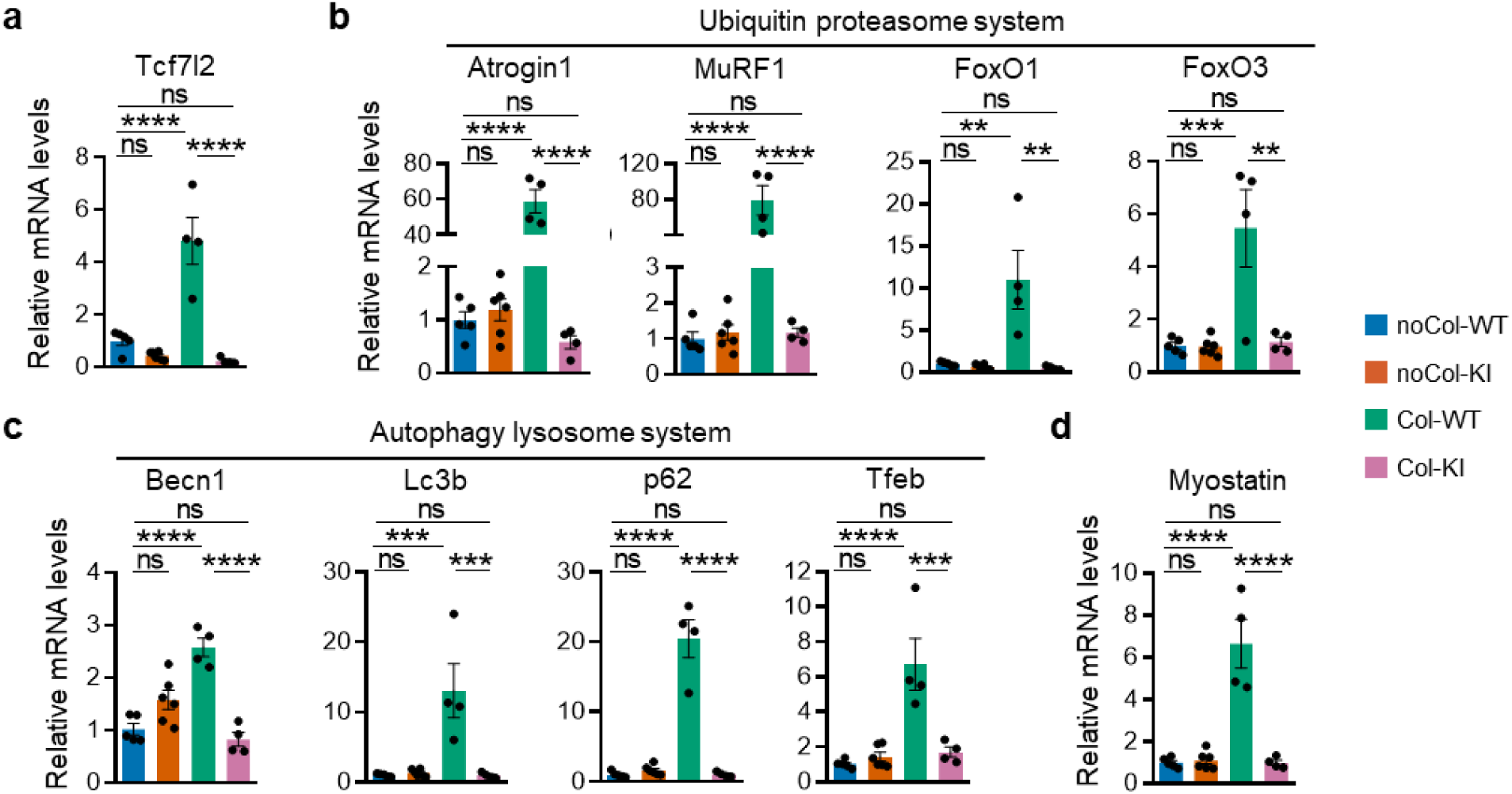
TCF7L2 regulates the expression of numerous atrophy genes in the gastrocnemius muscle. **(a-d)** Relative mRNA expression of **(a)** Tcf7l2, **(b)** ubiquitin-proteasome system genes, **(c)** autophagy-lysosome system genes and **(d)** myostatin in the gastrocnemius. Data are mean ± SEM. Statistical significance was assessed one-way ANOVA with Bonferroni test. noCol-WT *n* = 5, noCol-KI *n* = 6, Col-WT *n* = 4, Col-KI *n* = 4. **P < 0.01; ***P < 0.001; ****P < 0.0001 and ns, not significant.

Skeletal muscle atrophy results from protein degradation, which is governed mainly by the ubiquitin-proteasome system (UPS) and autophagy-lysosome system (ALS) in cancer cachexia (35). Hence, we measured the mRNA expression of key players in both systems. UPS genes, including *FoxO1* and *FoxO3* transcription factors (36), and *MuRF1* and *Atrogin1* ubiquitin ligases (37, 38) were highly upregulated in Col-WT gastrocnemius (Fig. 4b), which suggests that the UPS was hyperactivated. All four UPS genes were likely regulated by TCF7L2 because their expression returned to non-cachectic levels upon *Tcf7l2* repression in Col-KI gastrocnemius (Fig. 4b). ALS genes, including *Becn1*, *Lc3b*, *p62* and *Tfeb,* had similar hyperactivation profiles (Fig. 4c). As was the case for *Myostatin* (Fig. 4d), which induces the UPS via ActRIIB-Smad signaling (39) but independently of FOXO transcription factors. Taken together, these findings suggest that TCF7L2 is upregulated in the skeletal muscle during severe cachexia, and it promotes the expression of multiple atrophy genes to induce muscle loss. Hence, cachectic Col-KI mice recovered muscle mass loss upon *Tcf7l2* repression.

Since TCF7L2 is a transcription factor, we investigated whether it could bind to atrophy genes at the DNA level by using publicly available ChIP-Seq data from small intestine cells and liver hepatocytes (40, 41). Multiple TCF7L2 binding sites were present in the vicinity of all eight UPS and ALS genes (Supplementary Table S1); in particular, the MuRF1 enhancer and Atrogin1 promoter (Supplementary Fig. S5). While neither ChIP-Seq data were from skeletal muscles, this *in silico* analysis opens the possibility that TCF7L2 can act as a transcriptional activator to directly regulate the expression of atrophy genes in cachectic muscles.

*Tcf7l2* repression may have affected the expression of cachectic factors, to bring about cachexia recovery in Col-KI mice. To explore this possibility, we measured the mRNA levels of known cachectic proteins (17) and serum concentrations of known cachectic metabolites (18) in Col-WT and Col-KI mice. Although several cachectic proteins showed higher expression in tumors compared to their adjacent normal tissues, none were downregulated in Col-KI tumors (Supplementary Fig. S6a). Similarly, there was no difference in the concentrations of cachectic metabolites between Col-WT and Col-KI serum (Supplementary Fig. S6b). We then expanded our analysis to include GDF-15 (22–25) and the comprehensive set of 23 cachectic factors defined by *Friere et al.* (17), using our earlier bulk RNA-Seq data. However, only three cachectic factors differed significantly between Col-WT and Col-KI tumors, and all three were actually upregulated in Col-KI tumors (Supplementary Table S2). Therefore, TCF7L2 does not appear to regulate the expression of tumor-secreted cachectic factors.

### TCF7L2 is essential for early small intestine tumors

Next, we investigated whether TCF7L2 is required for small intestine tumors. Experimental and control mice had the same genotypes as colon cancer mice (Fig. 2a), but were labelled with the prefix “SI” and “noSI” instead of “Col” and “noCol” to differentiate between the two cancer models. DSS treatment was omitted since *Apc*^Min/+^ mice spontaneously develop small intestine tumors (42). When doxycycline was initiated early at 6 weeks of age (Fig. 5a-b) and before adenoma propagation (43), SI-KI small intestines showed a dramatic reduction in the number of lesions (Fig. 5c-d). Furthermore, histological analysis revealed that these lesions were restricted to dysplastic structures that had not yet progressed to adenomas, unlike SI-WT small intestines (Fig. 5e). Conversely, when doxycycline treatment was delayed until the onset of weight loss between 4 and 6 months of age (Fig. 6a-b), the tumor burden in SI-KI mice was unaffected (Fig. 6c-d) and nearly all lesions were adenomas (Fig. 6e). A check of *Tcf7l2* mRNA levels revealed only partial 60-80% repression in small intestine tumors (Fig. 6f), which translated to a slightly diminished expression of Wnt target genes (Supplementary Fig. S7). Overall, these results suggest that TCF7L2 is crucial for early tumourigenesis in the small intestine and even moderate repression of the gene can halt progression to adenomas. However, TCF7L2’s role in established small intestine tumors is less conclusive, due to incomplete gene repression.

**Figure 5.**
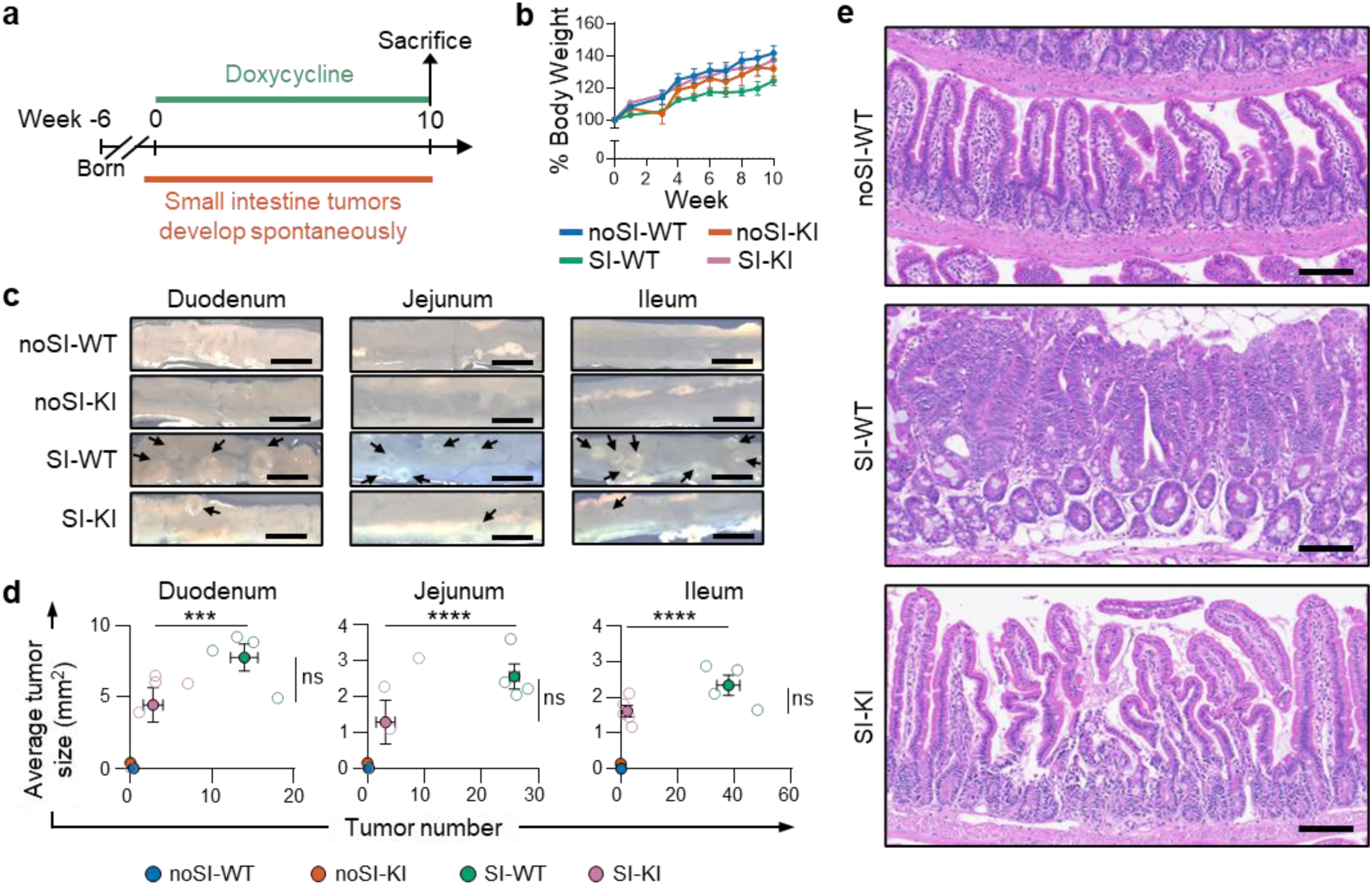
TCF7L2 is essential for early tumourigenesis in the small intestine. **(a)** Experimental timeline for small intestine tumor development and doxycycline treatment. **(b)** Body weight with respect to experimental day 0. **(c)** Representative photos of longitudinally opened small intestine. Arrows indicate tumor lesions. Scale bar 5 mm. **(d)** Quantification of tumor burden. **(e)** H&E staining of small intestine. Scale bar 100μm. Data are mean ± SEM. Statistical significance was assessed one-way ANOVA with Bonferroni test. noSI- WT *n* = 8, noSI-KI *n* = 4, SI-WT *n* = 4, SI-KI *n* = 3. ***P < 0.001; ****P < 0.0001 and ns, not significant.

**Figure 6.**
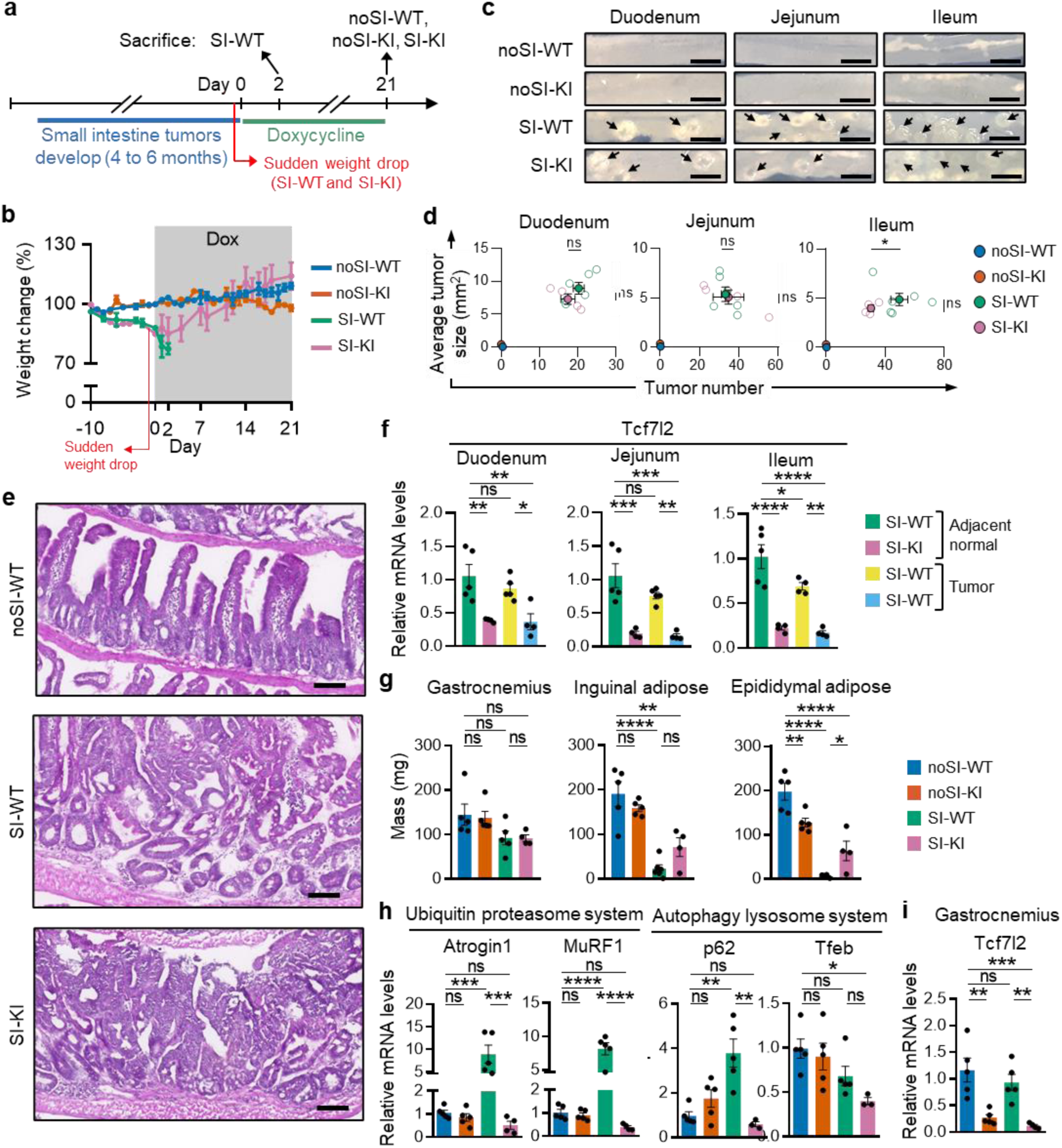
*Tcf7l2* repression rescues mice from fatal cachexia without affecting small intestine tumor growth. **(a)** Experimental timeline for tumor development and doxycycline treatment. **(b)** Body weight with respect to day-10. **(c)** Relative mRNA expression of *Tcf7l2* in tumors and the adjacent normal tissue. **(d)** Representative photos of longitudinally opened small intestine. Arrows indicate tumor lesions. Scale bar 5 mm. **(e)** Quantification of tumor burden. **(f)** H&E staining of small intestine. Scale bar 100μm. **(g)** Mass of the gastrocnemius and adipose tissues. **(h-i)** Relative mRNA expression of **(h)** Tcf7l2 and **(i)** muscle atrophy genes in the gastrocnemius. Data are mean ± SEM. Statistical significance was assessed one-way ANOVA with Bonferroni test. noSI-WT n = 5, noSI-KI n = 5, SI-WT n = 5, SI-KI n = 4. *P < 0.05; **P < 0.01; ***P < 0.001; ****P < 0.0001 and ns, not significant.

### *Tcf7l2* repression reverses fatal small intestine cancer cachexia

The onset of severe refractory cachexia in SI-WT and SI-KI mice between 4 and 6 months of age was marked by sudden body weight loss (Fig. 6b) and a drop in gastrocnemius, inguinal adipose and epididymal masses (Fig. 6g), similar to Col-WT and Col-KI mice. However, unlike the massive upregulation of atrophy genes in Col-WT gastrocnemius that reached as high as 80-fold (Fig. 4b-c), atrophy gene expression increased by a modest 15-fold or less in SI-WT gastrocnemius (Fig. 6h).

Despite this, doxycycline treatment reversed fatality (Fig. 6b) and significantly extended the lifespan of SI-KI mice by more than 10 weeks (Supplementary Fig. S8a-b). Unexpectedly, *Tcf7l2* was not elevated in the gastrocnemius of cachectic SI-WT mice (Fig. 5i) and gastrocnemius mass remained low in SI-KI mice when *Tcf7l2* was repressed (Fig. 6g), which suggests that recovery was independent of the gastrocnemius. Therefore, the results indicate that TCF7L2 drives severe cachexia in small intestine and colon cancer mice through distinct mechanisms. Interestingly, extending the lifespan of SI-KI mice through *Tcf7l2* repression led to the spontaneous development of numerous colon tumors (Supplementary Fig. S8a-c), a phenomenon that is not typically observed in *Apc*^Min/+^ mice (42).

## Discussion

Using a newly generated transgenic mouse to partially and ubiquitously repress *Tcf7l2*, we show that this gene is essential for the development of adenomas in the small intestine but not colon. Additionally, *Tcf7l2* is not required for the maintenance of both established small intestine and colon adenomas. More importantly, we discover an unprecedented link between TCF7L2 and cancer cachexia. In colon cancer cachexia, TCF7L2 directly drives muscle atrophy by regulating the expression of atrophy genes in the gastrocnemius muscle. In contrast, its role in small intestine cancer cachexia appears to be independent of atrophy gene regulation in the gastrocnemius. Overall, our findings demonstrate that both small intestine and colon cancer mice can be rescued from cachexia-induced death by repressing *Tcf7l2*.

Loss of the wildtype *Apc* allele in *Apc*^min/+^ mouse, which leads to hyperactivation of the canonical Wnt signaling pathway, is sufficient for adenoma formation in the small intestine (44, 45).

However, this typically only causes dysplasia in the colon (45), with additional mutations and epigenetic changes required for progression to adenomas (46). This difference in early tumor development between the two organs is underscored by our observation that *Tcf7l2* repression impacted adenoma development in the small intestine, but not in the colon. Since all lesions in the small intestine were dysplastic, it is possible that *Tcf7l2* repression either arrested tumor transformation at this stage or induced adenoma regression. Notably, this was achieved with only partial 60-80% *Tcf7l2* repression, implying that small intestine adenomas are highly dependent on TCF7L2 activity. Hence, even modest TCF7L2 inhibition might be sufficient for the preventive treatment of small intestine cancer in patients with Familial Adenomatous Polyposis (FAP).

Once small intestine and colon adenomas were established, they continued growing despite *Tcf7l2* repression. This contradicts the observation of Dow *et al.*’s (30) that both small intestine and colon tumors regress when the same hyperactivated Wnt signaling pathway is attenuated. A plausible explanation for the persistence of small intestine adenomas is that *Tcf7l2* was only partially repressed, so residual expression may have been adequate for tumor maintenance. However, this does not apply to colon adenomas, where repression exceeded 95%, and previous study has shown that colorectal cancer cell lines can survive biallelic *Tcf7l2* inactivation (8). Instead, the discrepancy might arise because we targeted different proteins along the Wnt signaling pathway. By stabilizing upstream *Apc*, Dow *et al.* (30) brought Wnt pathway activation back to pre-tumor levels. In contrast, the pathway remained slightly hyperactivated when we repressed downstream *Tcf7l2*, likely because the other three TCF/LEF proteins could transduce Wnt signal. In fact, all four TCF/LEF proteins regulate distinct aspects of colon tumor biology, as revealed by a meta-analysis using microarray data from colorectal cancer patients (47). TCF7L1, TCF7, and LEF1 are linked to angiogenesis, metabolic processes, and tumor migration, respectively (47). TCF7L2 is important for colorectal cancer cell proliferation, as gene disruption causes G1 cell cycle arrest via downregulation of the Wnt target gene c-Myc (48). Additionally, TCF7L2 may inhibit tumor metastasis, since knockout in colorectal cancer cell lines increases migration (8, 9). Our bulk RNA-Seq data corroborates these TCF7L2 roles; reduced expression of proliferation and Myc target genes but enrichment of EMT genes when *Tcf7l2* was repressed. Therefore, targeting TCF7L2 is unlikely to the reduce tumor burden in colorectal cancer patients because the gene is dispensable for tumor maintenance.

The *Apc*^Min/+^ mouse is an established model for cancer cachexia that is driven by small intestine tumors (49). In this model, weight loss and muscle atrophy occur gradually over several months (49) because the tumor burden closely mirrors human cancers. This is unlike tumor transplant models, where overtly large tumors can act as glucose and nitrogen traps (50, 51) to accelerate cachexia. As we demonstrate, the DSS-treated *Apc*^Min/+^ mouse also developed cachexia marked by gradual weight loss and muscle atrophy, eventually leading to death. In this model however, cachexia was driven by colon tumors rather than small intestine tumors, because symptoms appeared around three months of age—much earlier than small intestine cancer cachexia in untreated *Apc*^Min/+^ mice, which began between 4-6 months of age. Hence, the DSS-treated *Apc*^Min/+^ mouse can accurately model cancer cachexia and offers a unique perspective to the *Apc*^Min/+^ mouse since cachexia is driven by a different cancer type.

In both cancer cachexia models, we found that *Tcf7l2* repression led to a rapid and striking recovery of body weight, even in moribund cachectic mice. Although we could not determine exactly how *Tcf7l2* repression led to the recovery of small intestine cancer cachexia mice, the recovery mechanism in colon cancer cachexia mice was clearly tumor-extrinsic. This is because the expression of cachectic factors remained unchanged, and colon tumors persisted, sometimes progressing to invasive carcinomas. Instead, TCF7L2 behaved as a master regulator of multiple UPS and ALS atrophy genes in the skeletal muscle during colon cancer cachexia. Hence, its repression led to the downregulation of these atrophy genes and a recovery in muscle mass. A previous study showed that activation of the canonical Wnt signaling pathway in healthy fibro- adipogenic progenitors (FAPs), which are muscle stem cells, induces the secretion of ACTIVIN-A to promote atrophy gene expression and loss of volume in myofibers (52). It is possible that during cachexia, FAPs upregulate TCF7L2 to transduce Wnt signal and trigger this atrophy cascade.

However, since FAP numbers tend to decrease during cachexia (53), increased TCF7L2 expression may not fully compensate. Another possibility is that TCF7L2 is directly upregulated within myofibers, where it activates transcription of atrophy genes independently of FAPs. This direct mechanism would be consistent with the rapid onset and recovery that we observed in colon cancer cachexia mice.

Interestingly, extending the lifespan of *Apc*^Min/+^ mice by repressing *Tcf7l2* led to the spontaneous development of numerous tumors in the colon. Therefore, our transgenic mouse could be exploited for studying colon cancer by administering doxycycline early, before small intestine tumors emerge. This approach resembles sporadic colon cancer, unlike existing colitis-associated colon cancer models that induce tumors using DSS or azoxymethane (AOM). Since sporadic and colitis-associated colon cancers are driven by different molecular mechanisms (54), a sporadic colon cancer model could offer valuable insights into specific mechanisms and potential treatments.

A limitation of our study is that the colon tumors used for bulk RNA-Seq analysis were not age-matched. Therefore, while our interpretation of TCF7L2’s role in colon tumors aligns with that of other studies, it should be viewed with caution. A second limitation is that *Tcf7l2* was only partially repressed by 60-80% in small intestine tumors, due to concerns about lethality (3, 4, 6). Therefore, it was not possible to determine whether total repression would completely abolish early dysplastic structures and late established adenomas in the small intestine.

### Conclusions

In summary, our study provides a proof-of-concept that TCF7L2 could be targeted for therapeutic reversal of cachexia in small intestine and colon cancer patients. This has important clinical implications, particularly for refractory cachexia that is resistant to current treatment methods (12). TCF7L2’s role in muscle atrophy could potentially extend to aging sarcopenia (55), obesity (56) and neuromuscular disorders (57) since the Wnt pathway is involved in the pathogenesis of these muscle degenerative disorders. Finally, *TCF7L2* is the strongest risk locus associated with type II diabetes (58), so our transgenic mouse could be exploited for mechanistic studies of this disease.

## Methods

### Mice husbandry

All mouse procedures were performed according to the National Advisory Committee for Laboratory Animal Research (NACLAR) guidelines and approved by the Institutional Animal Care and Use Committee (IACUC) at Nanyang Technological University, Singapore (IACUC approval code AUP22074).

Mice were bred and housed under specific pathogen-free conditions under a 12h light/12h dark cycle. Maintenance diet and drinking water were provided, *ad libitum*. Experimental mice were of mixed genders and between 4-9 weeks old when treatment was initiated.

### Generation of transgenic mice

The Rosa26-rTetR-KRAB and Tcf7l2-TRE mouse strains were generated in-house by CRISPR- Cas9 targeting of embryonic stem cells (ESCs) (59–61) followed by blastocyst microinjection using standard transgenesis technique (62–65). Briefly, vectors were assembled as described below, and 1µg each of sgRNA and targeting vectors were transiently transfected into W4/129S6 ESCs using Fugene® HD (Promega corporation). ESCs clones carrying the targeted insertion were injected into C57BL/6 blastocysts, implanted into pseudopregnant females and carried to full-term. Transgene-positive male chimeric founders of both strains were back-crossed to C57BL/6 for two generations, before mating with *Apc*^Min/+^ mice. The resulting triple transgenic strain was maintained by inbreeding.

For the sgRNA vectors, sgRNA sequences specific for either the *Rosa26* locus (5’- GACTGGAGTTGCAGATCACG-3’) or *Tcf7l2* locus (5’-GTGCGTCTTGGGCTTTCCCC-3’), were cloned into the pX330-U6-Chimeric_BB-CBh-hSpCas9 vector (a gift from Feng Zhang; Addgene plasmid #42230), as described by Cong *et al.* (61).

For the Rosa26-rTetR-KRAB targeting vector, the expression cassette consisted of CAG promoter- rTetR-NLS-Kid1-poly(A). The rTetR sequence without VP16 domain was cloned from the rtTA vector by Gossen *et al.* (66), while partial Kid-1 sequence containing the minimal KRAB domain (amino acids 12 to 53) (67) was derived by reverse transcriptase-PCR from mouse tissue. The expression cassette was flanked by 5’ and 3’ homology arms corresponding to the *Rosa26* locus.

For the Tcf7l2-TRE targeting vector, the TRE_mod_ sequence from pTREtight vector (Clonetech) was flanked by 5’ and 3’ homology arms corresponding to the *Tcf7l2* locus. All homology arms were amplified from W4/129S6 ESCs genomic DNA.

### Genotyping

Tail clips were dissolved in SNET buffer pH8 (1% SDS, 400mM NaCl, 5mM EDTA, 20mM Tris-Cl, 200μg/ml proteinase K) at 55°C, and DNA was purified by ethanol precipitation. Purified DNA was amplified by multiplex PCR with the primers listed in Table S3. PCR products were separated on a 1% agarose gel containing ethidium bromide and visualized at 365nm.

### DSS and doxycycline treatment

Colon tumors were induced in *Apc*^Min/+^ mice by giving 1.5-2.5% dextran sodium sulphate (DSS) in the drinking water for 7 days, *ad libitum*. DSS treatment caused variable body weight loss of 5- 20%. *Tcf7l2* was repressed by feeding mice food pellets containing either 100mg/kg (partial repression) or 625mg/kg (complete repression) doxycycline hyclate, *ad libitum*. The base for doxycycline pellets was Teklad global 2018 diet (Envigo).

### Tissue harvesting

Mice were sacrificed by CO_2_ asphyxiation before harvesting various organs. Colons and small intestines were opened longitudinally and washed with ice-cold PBS. Tumors were pinched from underlying muscle layer using forceps, while adjacent normal tissues were sampled from macroscopically normal-looking areas. Tissues were then homogenized in TRIzol (Invitrogen) with an X360 electric homogenizer (Ingenieurbüro CAT). All other organs were also harvested and homogenized in TRIzol.

### RNA isolation and reverse-transcriptase PCR

Total RNA was purified from TRIzol homogenates by chloroform phase separation combined with spin column extraction. Purified RNA was reverse transcribed into single-stranded cDNA with oligo dT primers, and used for endpoint, semi-quantitative or quantitative PCR as described below.

End-point and Semi-quantitative PCR were performed using the primers listed in Table S4. Reaction cycle was repeated 40 times for end-point PCR and as indicated in the figures for semi-quantitative PCR. Extension time was 30s. PCR products were separated on a 1% agarose gel containing ethidium bromide and visualized at 365nm.

Quantitative PCR was performed on the Eco Real-Time PCR System (Illumina) using PrecisionFAST qPCR Master Mix with SYBR Green (Primerdesign Ltd) and the primers listed in Table S5. Reactions were performed in duplicate with 60°C annealing temperature and 30s extension. Cycle thresholds were determined automatically by the Eco Real-Time PCR software (Illumina), while relative expressions were calculated with the 2^-ΔΔCT^ method by normalizing to *β- actin*.

### Histology

Small intestines and colons were prepared according to Bialkowska *et al.* (68), except overnight fixation was performed with 4% paraformaldehyde at 4°C. Tissues were frozen in Tissue-Tek O.C.T. compound (Sakura Finetek), cryo-sectioned at 7µm thickness, transferred to Polysine slides, and stained as described below. An Eclipse 80i microscope system (Nikon) was used for visualization.

H&E staining was performed with Gill’s III hematoxylin solution, acid-alcohol destain, Scott’s reagent (83mM magnesium sulphate, 8mM sodium bicarbonate in H_2_O) and 0.5% alcoholic eosin Y. Sections were dehydrated and mounted with DPX mountant (Sigma-Aldrich).

For immunostaining, sections were heated at 95-98°C in citrate buffer pH 6.0, 20 min. The sections were then incubated in 0.1% Triton-X for 10min, 0.6% hydrogen peroxide for 15min, blocking solution (10% FBS in PBS) for 3h, rabbit anti-Tcf7l2 antibody (clone C48H11, Cell Signaling Technology; 1:50 dilution) for 48h at 4°C and HRP polymer-anti-rabbit IgG antibody (Dako) for 1h. Sections were visualized with 3,30-diaminobenzidine tetrahydrochloride (DAB), counterstained with Mayer’s hematoxylin, dehydrated and mounted with DPX mountant.

### Bulk RNA sequencing

RNA was purified from tumors as described above, but with an on-column DNase I digest. Total RNA samples with RIN >8 were converted to cDNA libraries and sequenced on HiSeq4000 (Illumina).

Paired-end reads were mapped to the Mouse GRCm38/mm10 reference genome using STAR (69) alignment and differentially expressed genes were identified with DESeq2 (70). Gene Set Expression Analysis was performed using Hallmark gene sets (70–72).

### Metabolomics analysis

Sera were prepared from whole blood and spiked with an internal standard, before extracting metabolites via cold methanol extraction. Samples were injected onto a UPLC HSS column (Waters Corporation) with water plus 0.1% formic acid and acetonitrile plus 0.1% formic acid as the mobile phases. Subsequently, mass spectrometry analysis was performed with an ESI positive and negative 6500+ QTRAP (SCIEX) and data was processed with the OS-MQ software (SCIEX). Metabolite concentrations were calculated by normalizing to the internal standard.

### Statistical analysis

Data were analyzed and plotted with GraphPad Prism 6.0 software. Statistical significance was determined using Student’s t-test or one-way ANOVA with Bonferroni correction, as stated in the figure legends.

## Declarations

### Data availability

The bulk RNA-Seq data generated in this study are publicly available in Gene Expression Omnibus (GEO) at GSE212581 (https://www.ncbi.nlm.nih.gov/geo/query/acc.cgi?acc=GSE212581). All other raw data are available at DR-NTU (https://doi.org/10.21979/N9/85ZLCG).

### Competing interests

The authors declare no potential conflicts of interest.

## Funding

This work was supported by the Ministry of Education Tier 1 grants (RG139/18 and RG22/22) awarded to C.R. and K.K..

## Authors’ contributions

M.L.L designed and executed the experiments, interpreted the results, and wrote the manuscript.

C.R and K.K supervised the project and discussed the results.

## Acknowledgements

We thank Dr Piotr Tetlak for assistance with establishing the transgenic strains, Dr Arun Kumar Manickam for advice on blastocyst injection and stock of mouse embryonic feeder cells and Dr Zhihao Wu for cloning the Rosa26-rTetR-KRAB targeting vector. RNA-Sequencing, alignment and differential gene expression analysis services were provided by the Genome Institute of Singapore. Metabolomics service was provided by the Singapore Phenome Centre.

## Notes

### Competing Interest Statement

The authors have declared no competing interest.

